# 16S rRNA Amplicon Sequencing of Bagworm *Metisa plana* Walker (Lepidoptera: Psychidae)

**DOI:** 10.1101/2021.03.30.437633

**Authors:** Andrew Chung Jie Ting, Cik Mohd Rizuan Zainal Abidin, Noor Hisham Hamid, Ghows Azzam, Hasber Salim

## Abstract

The bagworm *Metisa plana* is one of the major pests in the oil palm plantation in Malaysia, with infestation that results in huge economical loss. Currently, the exact cause of the infestation is still undetermined. Studying the bacterial community of *M. plana* could provide insight on the problem as the bacteria associated with insects often provide numerous benefits to the insect itself. Using 16S rRNA amplicon sequencing, the study was conducted to compare the composition of the bacterial communities of two larval stages (early instar stage and late instar stage) from outbreak area, as well as comparing the late instar stage larvae from non-outbreak and outbreak areas. Generally, the bacterial community was dominated by *Proteobacteria* and *Actinobacteria* phyla while the *Enterobacteriaceae* was found to be the dominant family. When comparing between the early and late instar stage, *Proteobacteria* phylum was found to be more abundant in the late instar stage (82.36%) than in the early instar stage (82.28%). At the family level, the *Enterobacteriaceae* was slightly more abundant in late instar stage (75.46%) than in early instar stage (75.29%). The instar stage was observed to have no significant impact on the bacterial variability and showed similar bacterial community structure. When comparing between the non-outbreak area and outbreak, *Proteobacteria* was significantly more abundant in the outbreak area (82.02%) than in the non-outbreak area (20.57%). However, *Actinobacteria* was significantly more abundant in the non-outbreak area (76.29%) than in the outbreak area (14.16%). At the family level, *Enterobacteriaceae* was more abundant in outbreak area (75.41%) than in non-outbreak area (11.67%). *Microbacteriaceae* was observed to be more abundant in the non-outbreak area (70.87%) than in the outbreak area (12.47%). Although the result showed no significant difference in bacterial variability between different areas, it the bacterial community structure was significantly different.

## Introduction

The Lepidoptera is a vastly diverse insect order, with many species considered as major pests of agricultural importance (1). The Lepidopteran pest bagworm is the most serious and economically important pests in the oil palm plantations in Malaysia (2–6). The bagworm outbreak can result in a terrible yield loss which can translate into millions of Ringgit Malaysia (Malaysia’s local currency) (4,7). Of the common species of bagworm found in the oil plantations (*Mahasena corbetti, Pteroma pendula*, and *Metisa plana*), the *M. plana* is the most serious leaf defoliator (5,8,9). Although there are available and effective control measures (4,10–12), the outbreak and infestation of the bagworm is still an occurring problem due to the lack of understanding of the pests (2,3).

Huge ranges of microorganisms colonize the insects, from the largest of fungi to the smallest of virus. The microbiota composition of the insects differs greatly and are affected by different factors such as insect developmental stages, environments, and even diet (13–16). Often times, these microorganisms provide various benefits to the wellbeing of the insect, but sometimes may be pathogenic (13,17–19). An example of benefits from insect-bacteria interaction is the acquisition of nutrients. Chewing insects that feed on leaves would not have enough nitrogen solely from their diet. This insufficient nitrogen obtained from the diet would be supplemented by bacterial symbionts which can fix nitrogen and convert it into appropriate nitrogen-containing compounds (13,20,21). Some symbiotic bacteria could also protect the host against pathogens. In a separate study, the authors showed that the dominant symbiotic bacterium *Enterococcus mundtii* actively secretes bacteriocin against bacterial invaders. This interaction protects the host from other invading bacteria and at the same time, provides the bacterium an advantage which contributed to its dominance (22).

The bacterial community of the *M. plana* bagworm to the best of the author’s knowledge has yet to be explored. The current study therefore aims at identifying and compare the bacterial community of the insect host which could provide an insight to the cause of the outbreak. This knowledge can potentially be used to improve on the integrated pest management methods such as using microbes as a biocontrol agent (23–26). Here in this study, we used 16S rRNA amplicon sequencing to access and compare the bacterial community of the early instar stage and late instar stage larvae of bagworm *M. plana*. The study also accessed and compared the bacterial community of the larval *M. plana* from non-outbreak area as well as the outbreak area.

## Methods and Materials

### Ethic Statements

*Metisa plana* larvae of both early instar stage and late instar stage were collected from outbreak area located in Felda Gunung Besout 02/03, Trolak, Perak. The *M. plana* larvae of late instar stage was collected from non-outbreak area located in Felda Jengka 7, Jengka, Pahang. This species is a pest and is not protected by law. Bagworm was declared a dangerous pest under the Malaysia Act 167, Plant Quarantine Act 1976 (29). Sampling was performed with proper protective equipment to ensure no contamination from and to the bagworm samples.

### Total DNA Extraction

Genomic DNA (gDNA) was extracted using Qiagen DNeasy Blood and Tissue Kit with slight modifications (Cat No./ID: 69506) in 4 replicates. For each instar stage from outbreak area (early instar stage and late instar stage), 20 bagworms were removed from their bags and placed in 1.5 mL microcentrifuge tube before adding 180 µL of ATL buffer. The samples were then kept at −20 °C for 30 min before being homogenized using micropipette tips. Twenty microlitre of proteinase K was added to the sample and mixed by vortexing before the samples were incubated at 56 °C for 10 min. The samples were then vortexed for 15 sec before adding 200 µL of AL buffer. The samples were mixed by vortexing and incubated at 56 °C for 10 min. Ice-cold absolute ethanol of 200 µL was added to the samples and mixed. The samples were centrifuged at 6,000 × g for 1 min and the supernatant were transferred to DNeasy Mini spin column. The spin columns were then centrifuged at 6,000 × g for 1 min. The spin columns were placed in a new 2 mL collection tubes and 500 µL of Buffer AW1 was added before centrifuging for 1 min at 6,000 × g. The spin columns were again placed in new 2 mL collection tubes and added with 500 µL of Buffer AW2 before centrifuging at 13, 200 × g for 8 min. The spin columns were placed in new 1.5 mL microcentrifuge tubes and 50 µL of Buffer AE was added directly to the spin columns’ membranes. They were then incubated for 3 min at room temperature before centrifuging at 6, 000 × g for 1 min. The eluates were pipetted back into the spin column’s membrane and incubated for 3 min before centrifuging at 6,000 × g for 1 min. Gel electrophoresis was performed and the results were visualized under ultraviolet light. The DNA extraction was repeated using late instar stage larvae from non-outbreak area.

### Library Preparation and 16S Amplicon Sequencing

The extracted gDNA were sent to the sequencing service provider, Apical Scientific Sdn Bhd (https://apicalscientific.com/) for library preparation and sequencing. V3-V4 variable regions of the 16S ribosomal RNA gene was amplified using the forward primer (5’ CCTACGGGNGGCWGCAG) and reverse primer (5’ GACTACHVGGGTATCTAATCC).The selected regions were amplified again using locus specific sequence primers with overhang adapters (Table 1). After passing the quality check, the V3-V4 variable region were amplified using locus-specific sequence primers with overhang adapters (Table 1). All the PCR reactions were carried out with Q5® Hot Start High-Fidelity 2X Master Mix.

**Table 1.**
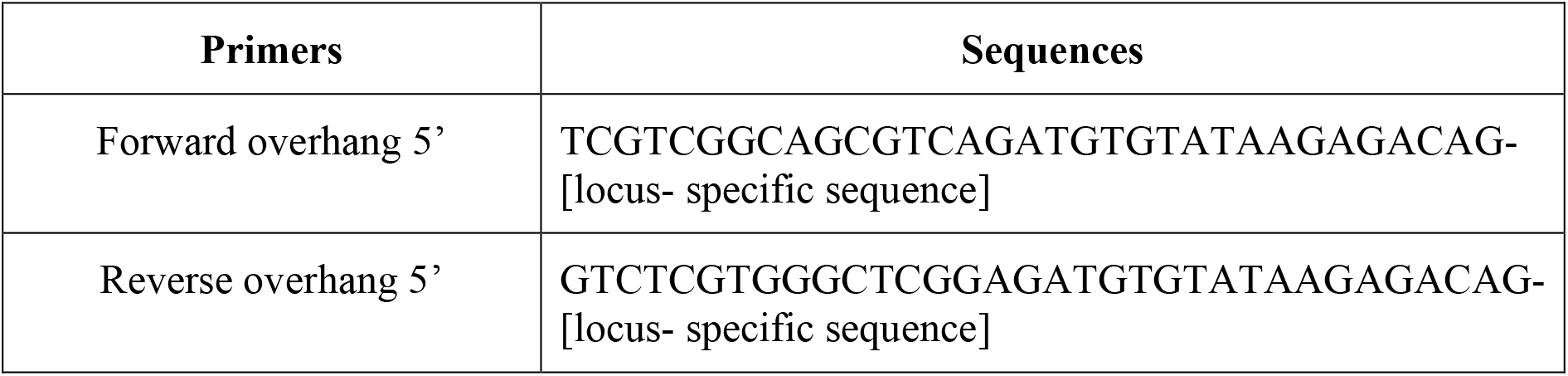
Overhang adapters used in library preparation.

## Analysis of Microbial Community

### Sequence Analysis

The analysis was done using Mothur software (v.1.44.3) with adaptations from MiSeq standard operating procedure (SOP) (https://mothur.org/wiki/miseq_sop/) (28). The forward reads and reverse reads were merged, and primers were removed. Sequences that were longer than 440 base pair (bp), but shorter than 406bp, and with any ambiguities were removed. Duplicates sequences and sequences that only appeared once were also removed. A customized reference targeting the V3-V4 region of the 16S rRNA gene was made from SILVA Seed v132 (29). Unique sequences were then aligned to the customized refence. Sequences that start before position 2 and ends after 17012, with homopolymer more than 8 as well as shorter than 406 bp were removed before removing gap characters. The sequences were pre-clustered, and chimeras were removed. The remaining sequences were classified to SILVA reference database using Bayesian classifier at 80% confidence threshold. Sequences that were classified into “Chloroplast”, “Mitochondria”, “Unknown”, “Archaea” and “Eukaryote” were removed. The sequences with similarity of 97% were then clustered into operational taxonomical units (OTU).

### Bacterial Community Analysis

As the samples showed unequal sampling depth, we investigated the alpha and beta diversity of the bacterial communities using rarefied OTU tables. To access the alpha-diversity, we calculated the Shannon diversity index, number of OTUs and Shannon evenness index. A simple T-test was performed to see whether the alpha diversity was significantly different. Principle Coordinate Analysis (PCoA) was plotted to visualise the cluster separation of the bacterial community’s structure. Analysis of Molecular Variance (AMOVA) was performed to see whether the centre of the cluster representing each group were significantly different. We performed Homogeneity of Molecular Variance (HOMOVA) to see whether the variation in each group were significantly different from each other.

## Results

### Overview of the Bacterial Community in *M. plana* larvae

From the overall results of this study, it was observed that the bacterial community of *M. plana* larvae was dominated by bacteria from the phyla *Proteobacteria* and *Actinobacteria*. At the family level, the bacterial community were generally dominated by *Enterobacteriaceae*. A detailed result of the comparison is explained systematically as follows.

### Comparison Between Early Instar and Late Instar Stage

#### Bacterial community composition of *M. plana* larvae

To obtain the bacterial community composition of the *M. plana* larvae at early instar and late instar stage, the V3 and V4 region of the bacterial 16S rRNA gene was amplified. A total of 2,738,727 sequences were obtained from 8 samples. After quality checks and removing unwanted sequences, a total of 385,297 sequences with 3,757 unique sequences were obtained. The sequences were then clustered at 97% similarity into 959 Operational Taxonomical Units (OTUs). The rarefaction curve did not completely plateau (Figure 1), suggesting the sequencing depth was insufficient to capture the entire bacterial community.

**Figure 1.**
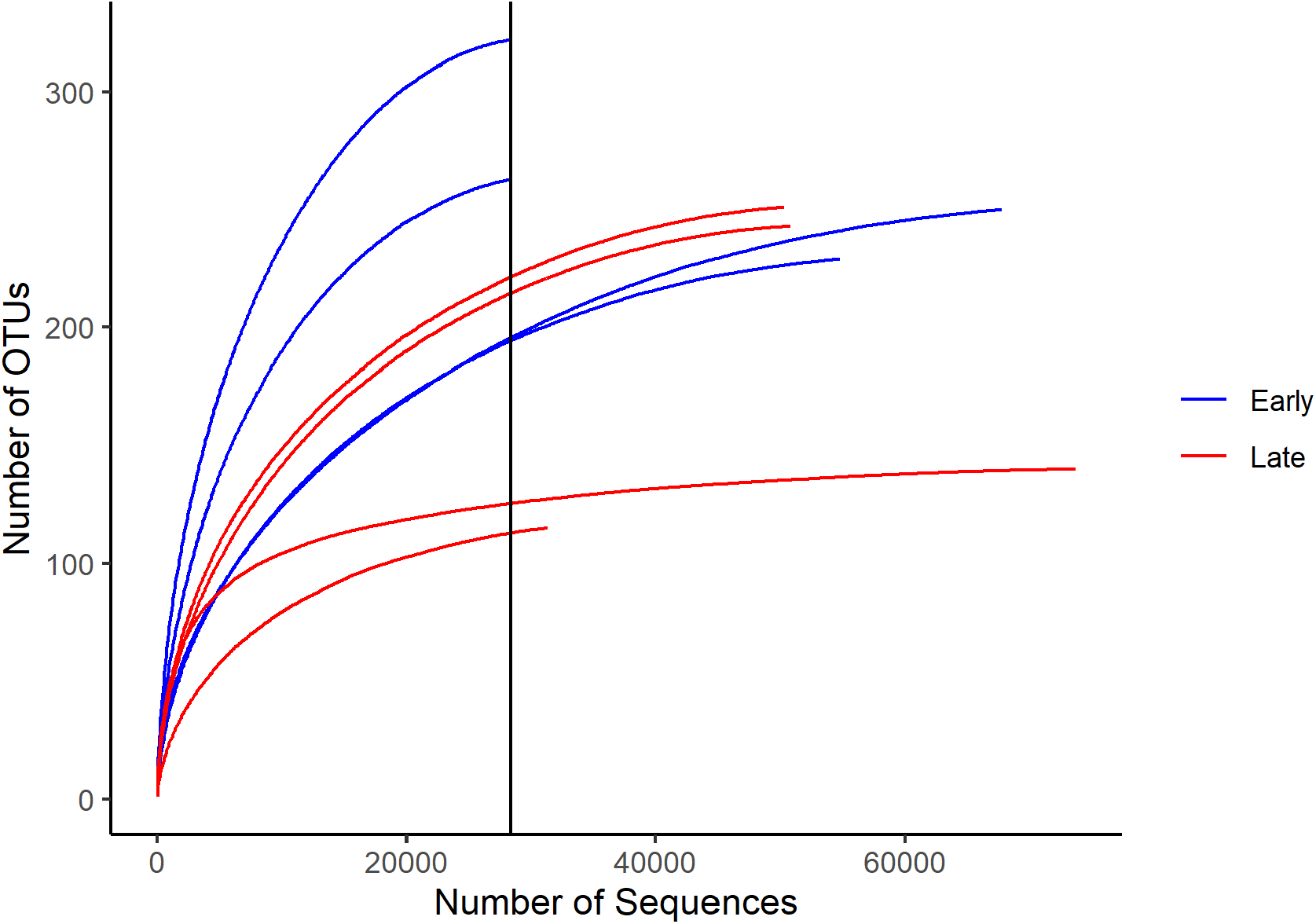
Rarefaction curve for the early instar stage and late instar stage samples. (x-axis intercept: samples were subsampled to 28,340 sequences). The curves showed that the early instar stage larvae generally have a higher number of OTUs.

#### Variability of bacterial communities between early instar stage and late instar stage

The bulk of the bacteria were of *Proteobacteria* (82.36%), *Actinobacteria* (14.8%), *Bacteroidetes* (1.48%), *Firmicutes* (1.01%) and remaining individual phyla consisting of less than 1% (Figure 2 and Table 2). However, there was no significant difference in relative abundance in any of the bacterial phyla.

**Table 2.**
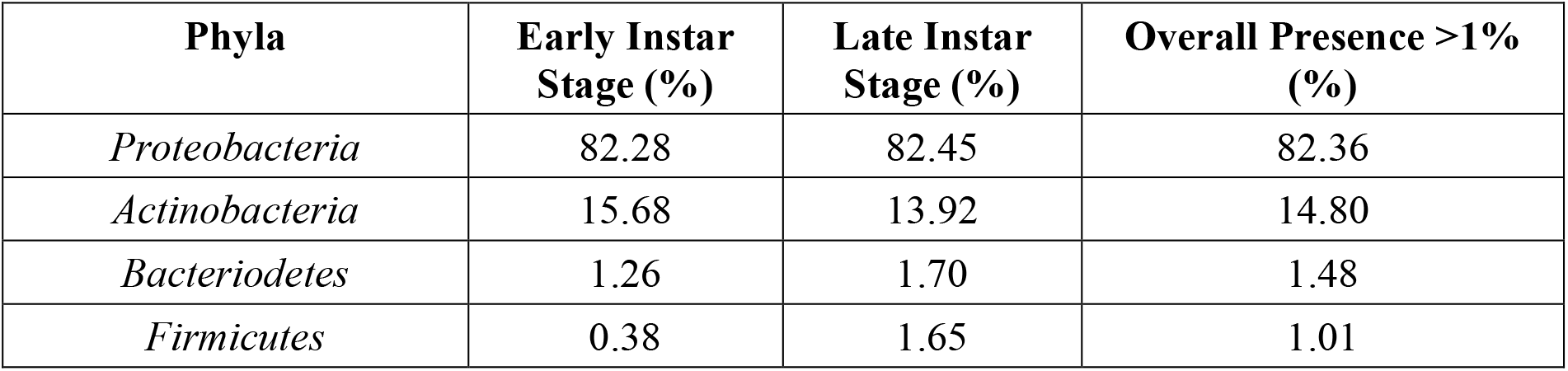
Bacterial phyla with an overall relative abundance of more than 1% in the comparison between early instar stage and late instar stage.

| Phyla | Early Instar Stage (%) | Late Instar Stage (%) | Overall Presence >1% (%) |
| --- | --- | --- | --- |
| *Proteobacteria* | 82.28 | 82.45 | 82.36 |
| *Actinobacteria* | 15.68 | 13.92 | 14.80 |
| *Bacteroidetes* | 1.26 | 1.70 | 1.48 |
| *Firmicutes* | 0.38 | 1.65 | 1.01 |

**Figure 2.**
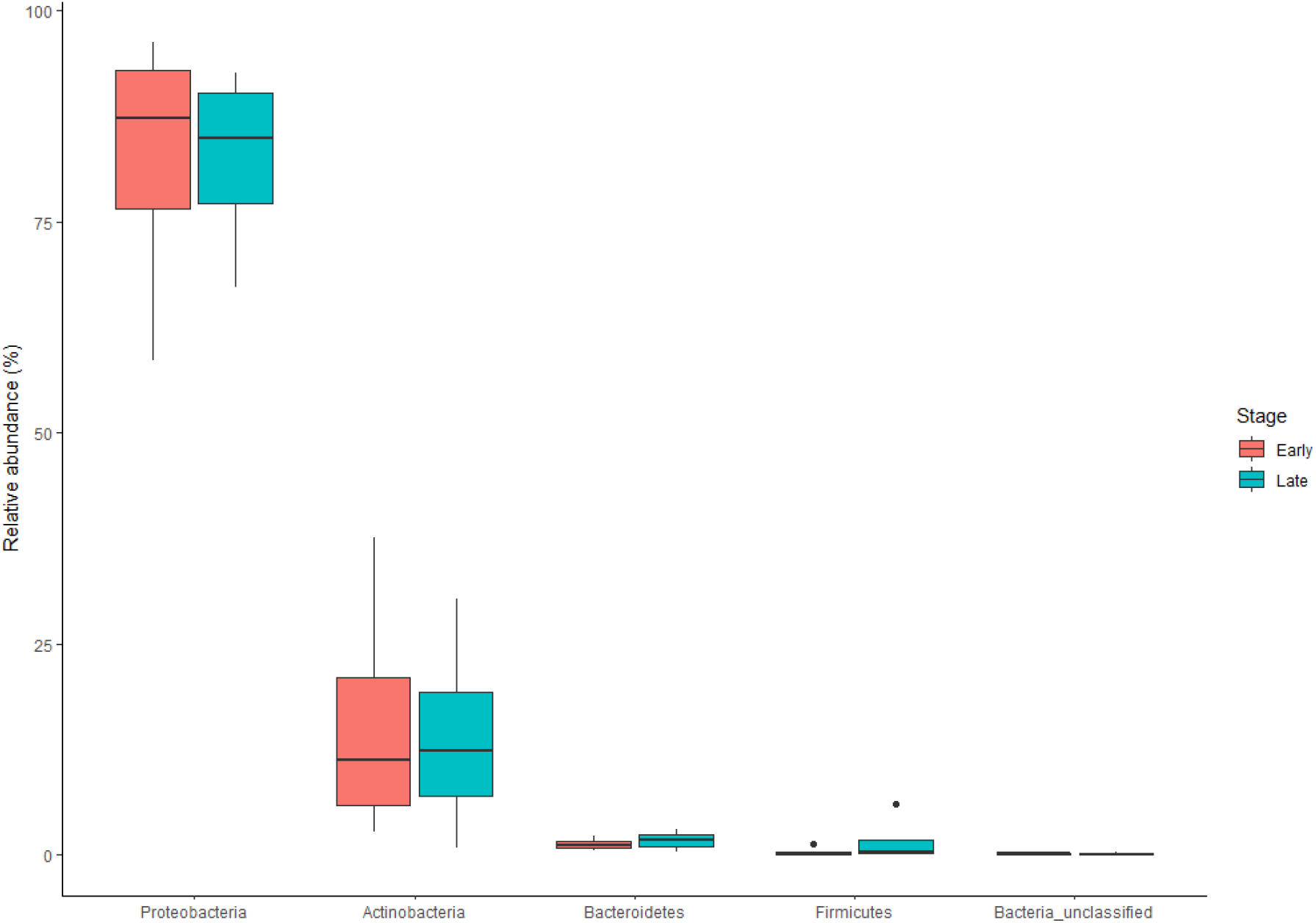
Top 5 relatively abundant bacterial phyla of *M. plana* bagworm larvae in the comparison between early instar stage and late instar stage.

At family level, the *Enterobacteriaceae* was the dominant family (75.37%), followed by *Microbacteriaceae* (13.63%), *Burkholderiaceae* (3.44%), *Pseudomonadaceae* (2.56%), *Sphingobacteriaceae* (1.09%) and the remaining families individually having less than 1% relative abundance (Figure 3 and Table 3). There was no significant difference in relative abundance between the dominant families, but there were a few minor families that were significantly differently in between the instar stage such as *Flavobacteriaceae, Legionellaceae, Nocardioidaceae* and *Pseudonocardiaceae*. There were more *Flavobacteriaceae, Nocardioidaceae* and *Pseudonocardiaceae* in the late instar stage, but the *Legionellaceae* was more abundant in the early instar stage (Table 4).

**Table 3.** Bacterial families with an overall relative abundance of more than 1% in the comparison between early instar stage and late instar stage.

| Families | Early Instar Stage (%) | Late Instar Stage (%) | Overall Presence >1% (%) |
| --- | --- | --- | --- |
| *Enterobacteriaceae* | 75.29 | 75.46 | 75.37 |
| *Microbacteriaceae* | 75.29 | 75.46 | 13.63 |
| *Burkholderiaceae* | 4.40 | 2.49 | 3.44 |
| *Pseudomonadaceae* | 1.59 | 3.53 | 2.56 |
| *Sphingobacteriaceae* | 0.99 | 1.18 | 1.09 |

**Table 4.** Bacteria with significant difference in relative abundance between early instar stage and late instar stage.

| **Families** | **Early instar (%)** | **Late instar (%)** | **Overall Presence (%)** |
| --- | --- | --- | --- |
| *Pseudonocardiaceae* | 0.04 | 0.28 | 0.16 |
| *Flavobacteriaceae* | 0.00 | 0.06 | 0.03 |
| *Nocardioidaceae* | 0.00 | 0.04 | 0.02 |
| *Legionellaceae* | 0.03 | 0.00 | 0.01 |

**Figure 3.**
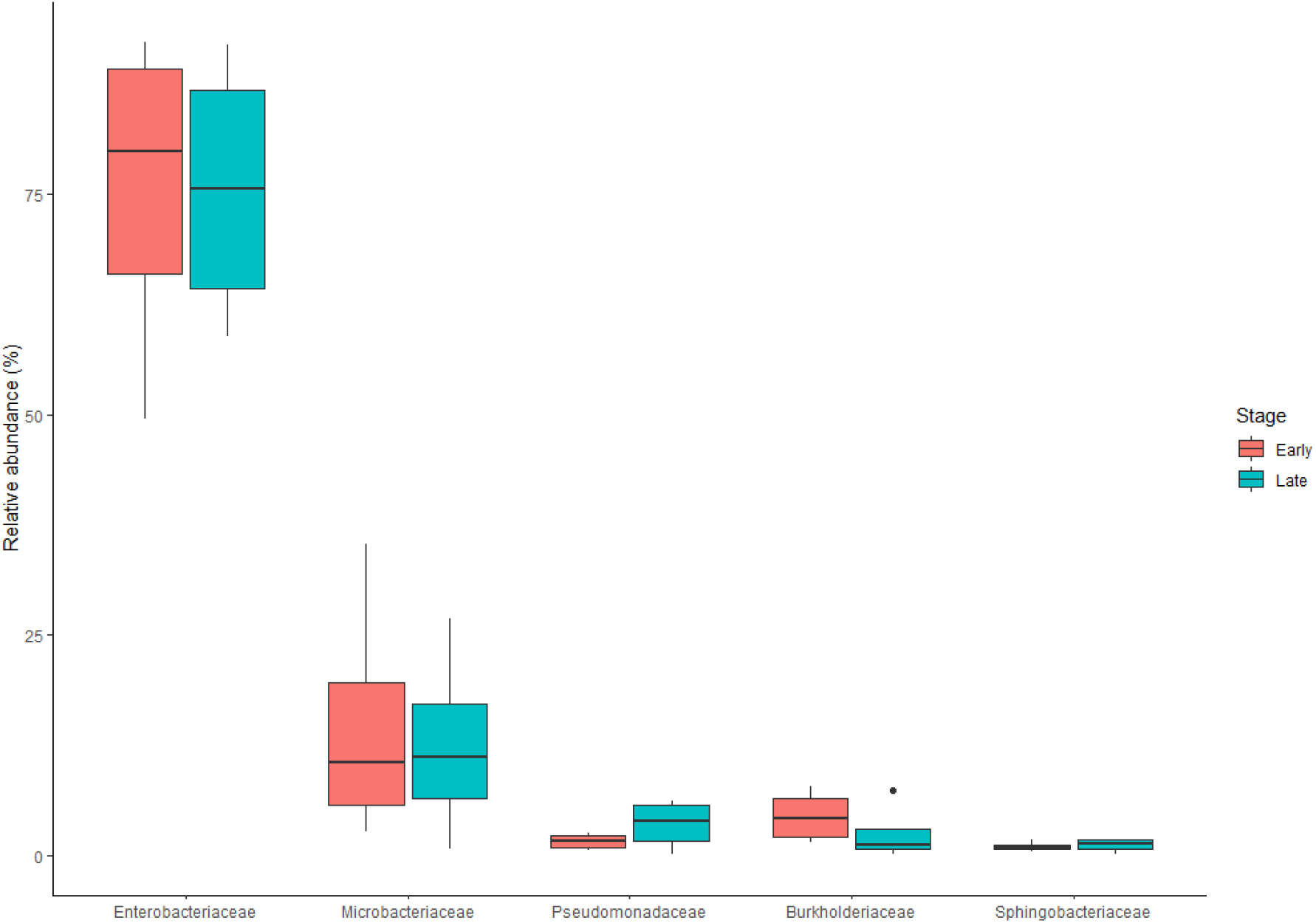
Top 5 relatively abundant bacterial families of *M. plana* bagworm larvae in the comparison between early instar stage and late instar stage.

#### Diversity of the bacterial community

Shannon diversity index were calculated to estimate the diversity of the bacterial community in the early and late instar stage (Figure 4 and Table 5). The index showed that the bacterial community of early instar stage was on average, more diverse than that of the late instar stage. The number of OTUs was also higher in the early instar stage than the late instar stage, revealing that the early instar stage was richer than the counterpart. Shannon evenness was obtained to observe the evenness of the bacterial community, and it showed that the bacterial community in early instar stage was more even than the late instar stage. However, the shannon diversity index, number of OTUs and evenness between the early instar stage and late instar stage were all not significantly different.

**Table 5.**
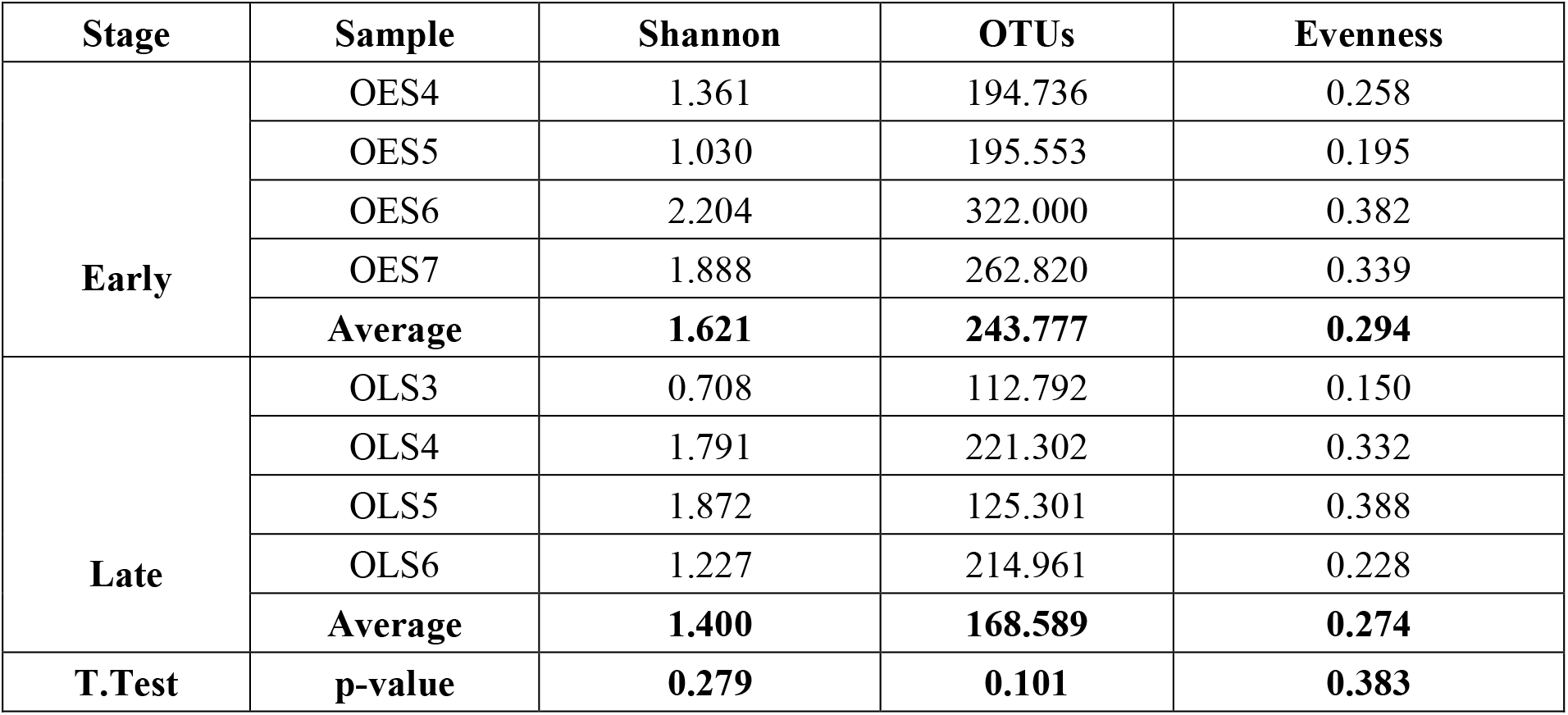
Shannon diversity index, Number of OTUs and Shannon evenness of bacterial community in the early instar and late instar stage. (Significance at p-value <0.05)

**Figure 4.**
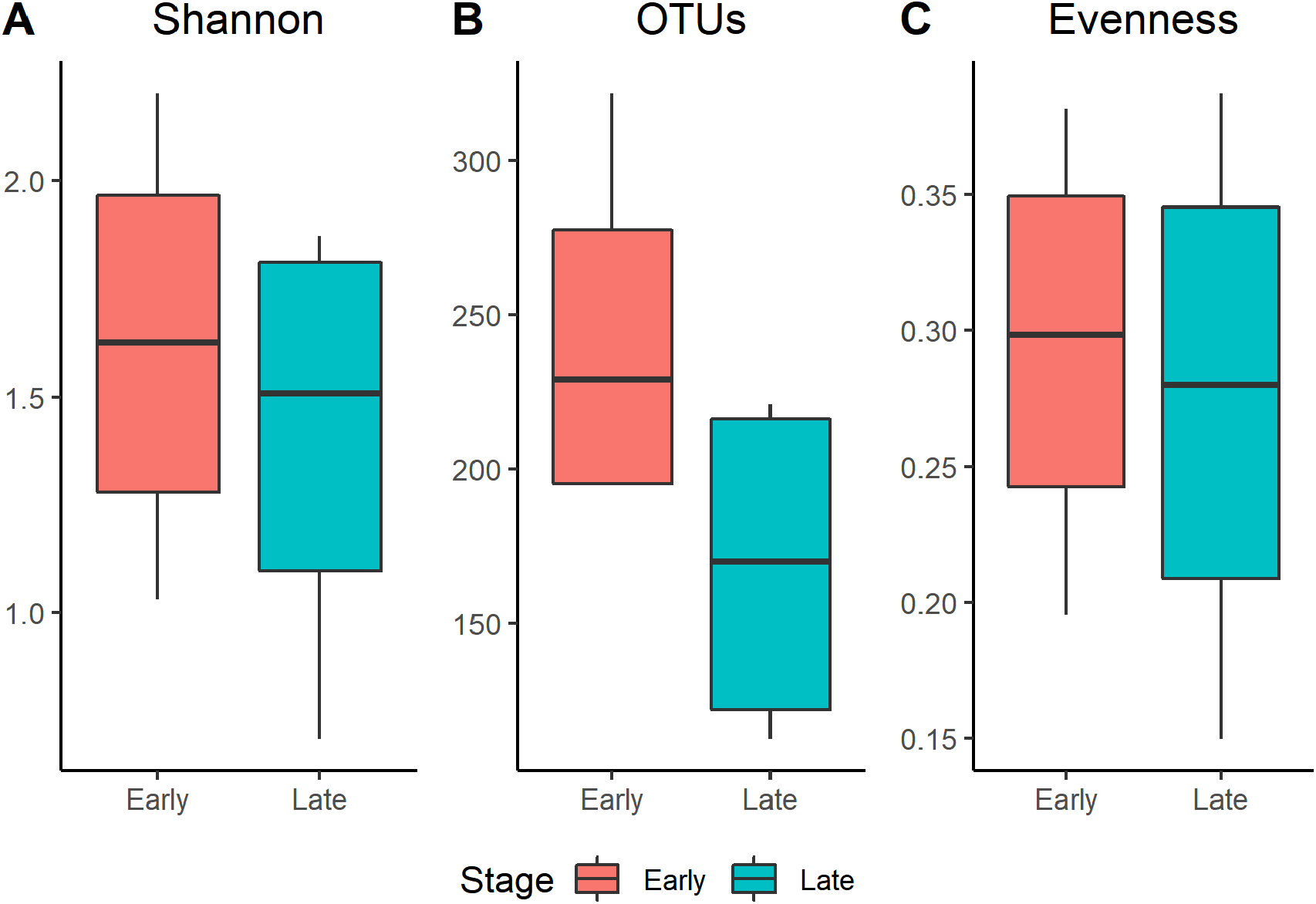
Alpha-diversity of the larvae of *M. plana* in comparison between instar stage. **A:** Shannon diversity index; **B:** Number of OTUs; **C:** Shannon Evenness

The PCoA was ordinated to visualise the cluster separation of the bacterial community. However, the ordination (Figure 5) did not show clear separation between the early instar stage and late instar stage. AMOVA test was done on the samples to test whether the cluster of the early instar and late instar stage was significantly different. The result of AMOVA (Table 6) revealed that the observed separation in the early instar and late instar stage was not significantly different. This meant that the bacterial community structure is like one another.

**Table 6.** AMOVA test done on samples from early instar stage and late instar stage. (Significance at p-value < 0.05)

| Early - Late | Among | Within | Total |
| --- | --- | --- | --- |
| SS | 0.010 | 0.191 | 0.201 |
| df | 1 | 6 | 7 |
| MS | 0.010 | 0.032 |  |
| Fs: | 0.325 |  |  |
| p-value: 0.554 |  |  |  |

**Figure 5.**
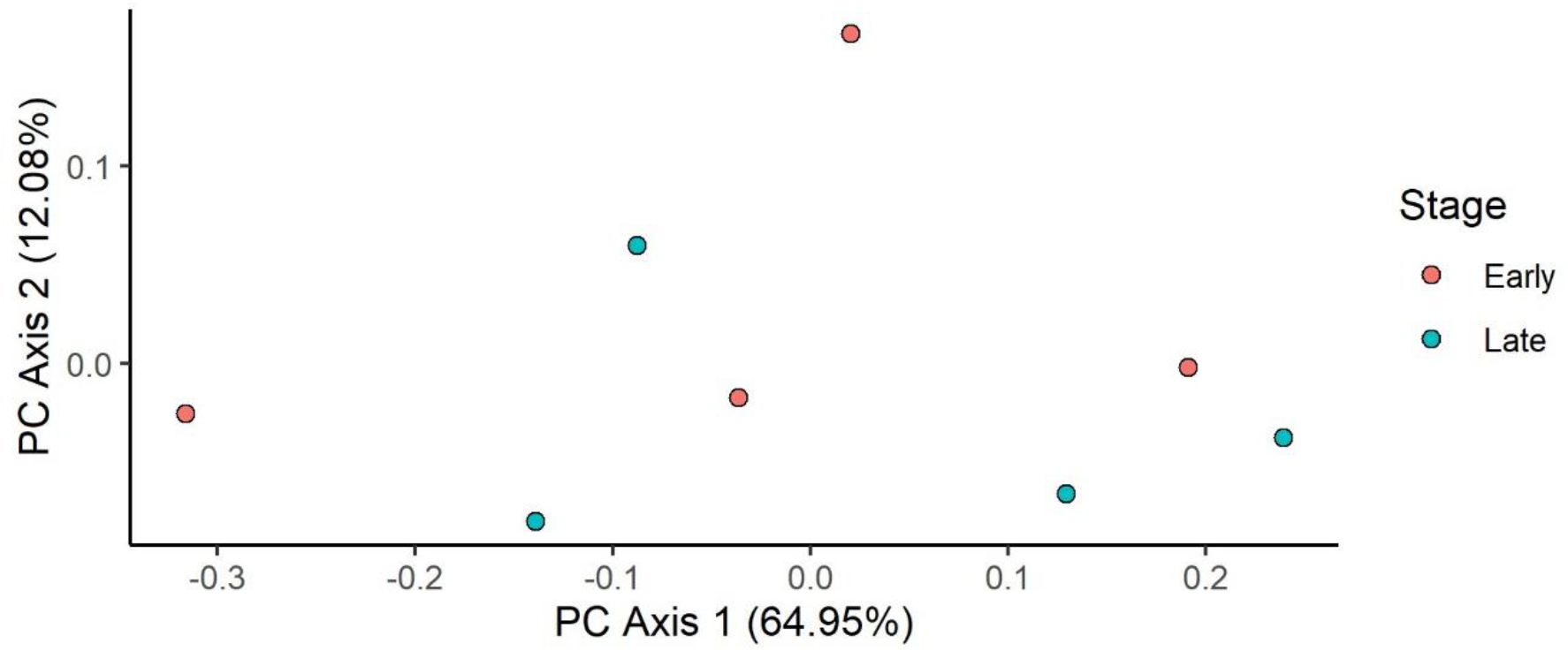
Principal Coordinate Analysis (PCOA) plot of bacterial communities of *M. plana* bagworm larvae in the comparison between early instar stage and late instar stage.

The authors also wanted to know whether the variation of the bacterial community in the early instar stage larvae was significantly different from that of the late instar stage. This was done by performing HOMOVA. From the HOMOVA test (Table 7), it showed that there was no significant difference in the variation with the early instar stage and late instar stage. Nonetheless, the early instar stage has a higher variation (0.038) compared to the late instar stage (0.026). This showed that bacterial community in the early instar stage was less stable than the late instar stage.

**Table 7.**
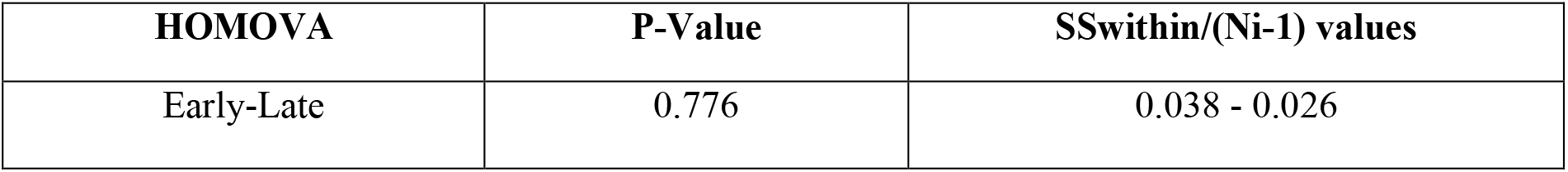
HOMOVA test done on the samples from early instar stage and late instar stage. (Significance at p-value<0.05)

| HOMOVA | P-Value | SSwithin/(Ni-1) values |
| --- | --- | --- |
| Early-Late | 0.776 | 0.038 - 0.026 |

### Comparison Between Non-Outbreak Area and Outbreak Area

#### Bacterial community composition of *M. plana* bagworm larvae from non-outbreak and outbreak area

The V3 and V4 region of the bacterial 16S rRNA gene was amplified using late instar stage larvae from the non-outbreak area and outbreak area. A total of 2,848,936 sequences were obtained from 8 samples. After quality checks and removing unwanted sequences, a total of 271,821 sequences with 2,471 unique sequences were obtained. The sequences were then clustered at 97% similarity into 796 Operational Taxonomical Units (OTUs). The rarefaction curve did not plateau (Figure 6), suggesting the sequencing depth was insufficient to capture the entire bacterial community.

**Figure 6.**
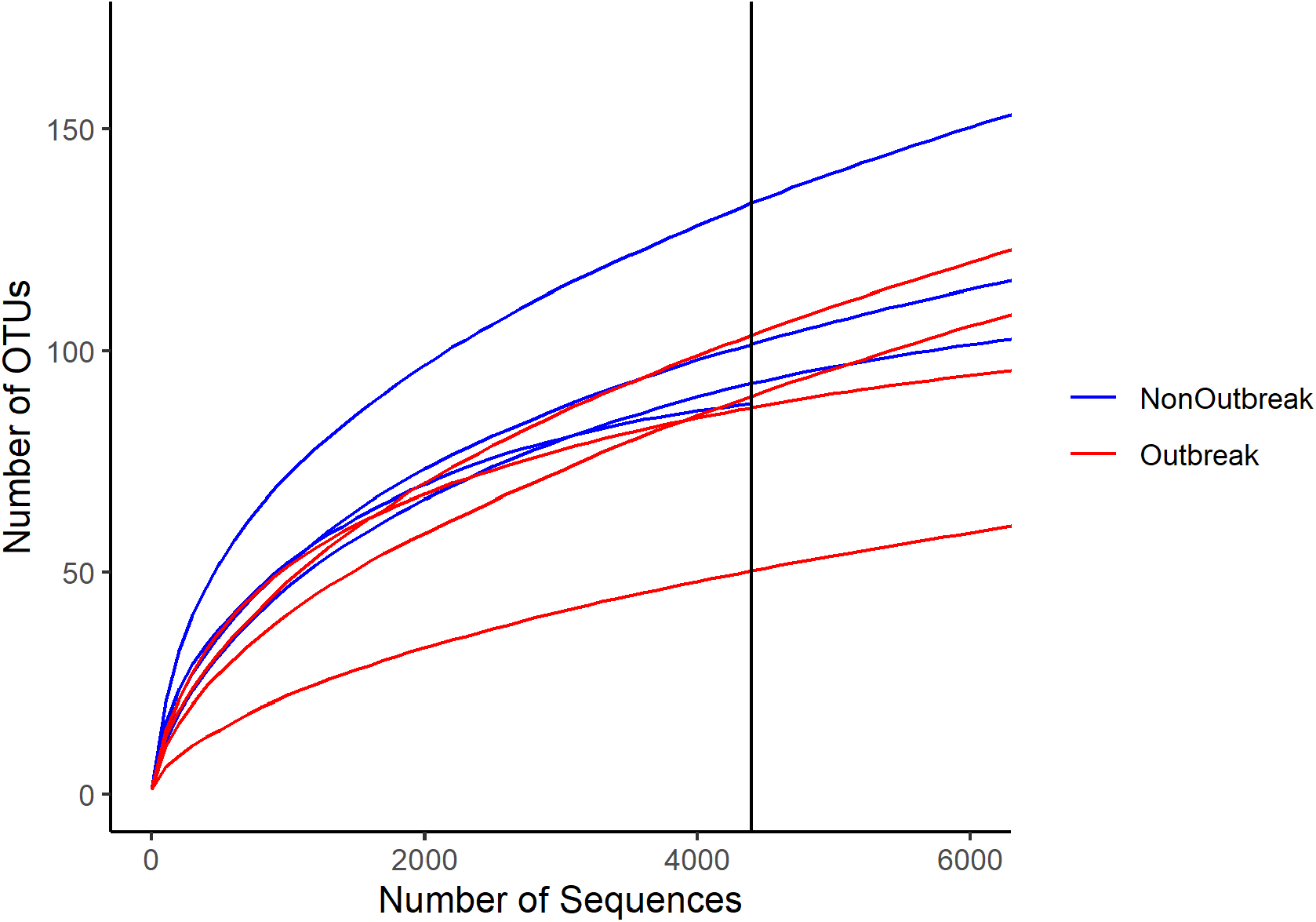
Rarefaction curve for the early instar stage and late instar stage samples. (x-axis intercept: samples were subsampled to 4,399 sequences). The curves showed the same number of sequences, the larvae from non-outbreak area had a greater number of OTUs than that of outbreak area.

#### Variability of bacterial communities between non-outbreak area and outbreak area

The most abundant phyla consisted of *Proteobacteria* (51.30%) followed by *Actinobacteria* (45.22%), *Bacteroidetes* (1.98%) and the rest of the phyla individually consisting of less than 1% in relative abundance (Figure 7 and Table 8). We observed a few phyla that were significantly different in relative abundance (Table 9). The *Proteobacteria* phylum from the outbreak area (82.02%) was greater in relative abundance than of non-outbreak area (20.57%). However, the second most dominant phylum which was the *Actinobacteria* was higher in relative abundance in non-outbreak area (76.29%) than that of outbreak area (14.16%). The unclassified bacteria in non-outbreak area (0.60%) were also greater than in outbreak area (0.14%).

**Table 8.** Bacterial families with an overall relative abundance of more than 1% in the comparison between non-outbreak area and outbreak area.

| Phyla | Non-outbreak area (%) | Outbreak area (%) | Overall Presence >1% (%) |
| --- | --- | --- | --- |
| *Proteobacteria* | 20.57 | 82.02 | 51.30 |
| *Actinobacteria* | 76.29 | 14.16 | 45.22 |
| *Bacterioidetes* | 2.19 | 1.76 | 1.98 |

**Table 9.** Bacteria with significant difference in relative abundance between non-outbreak area and outbreak area.

| Phyla | Non-outbreak area (%) | Outbreak area (%) | Overall Presence (%) |
| --- | --- | --- | --- |
| *Proteobacteria* | 20.57 | 82.02 | 51.30 |
| *Actinobacteria* | 76.29 | 14.16 | 45.22 |
| Unclassified bacteria | 0.60 | 0.14 | 0.37 |

**Figure 7.**
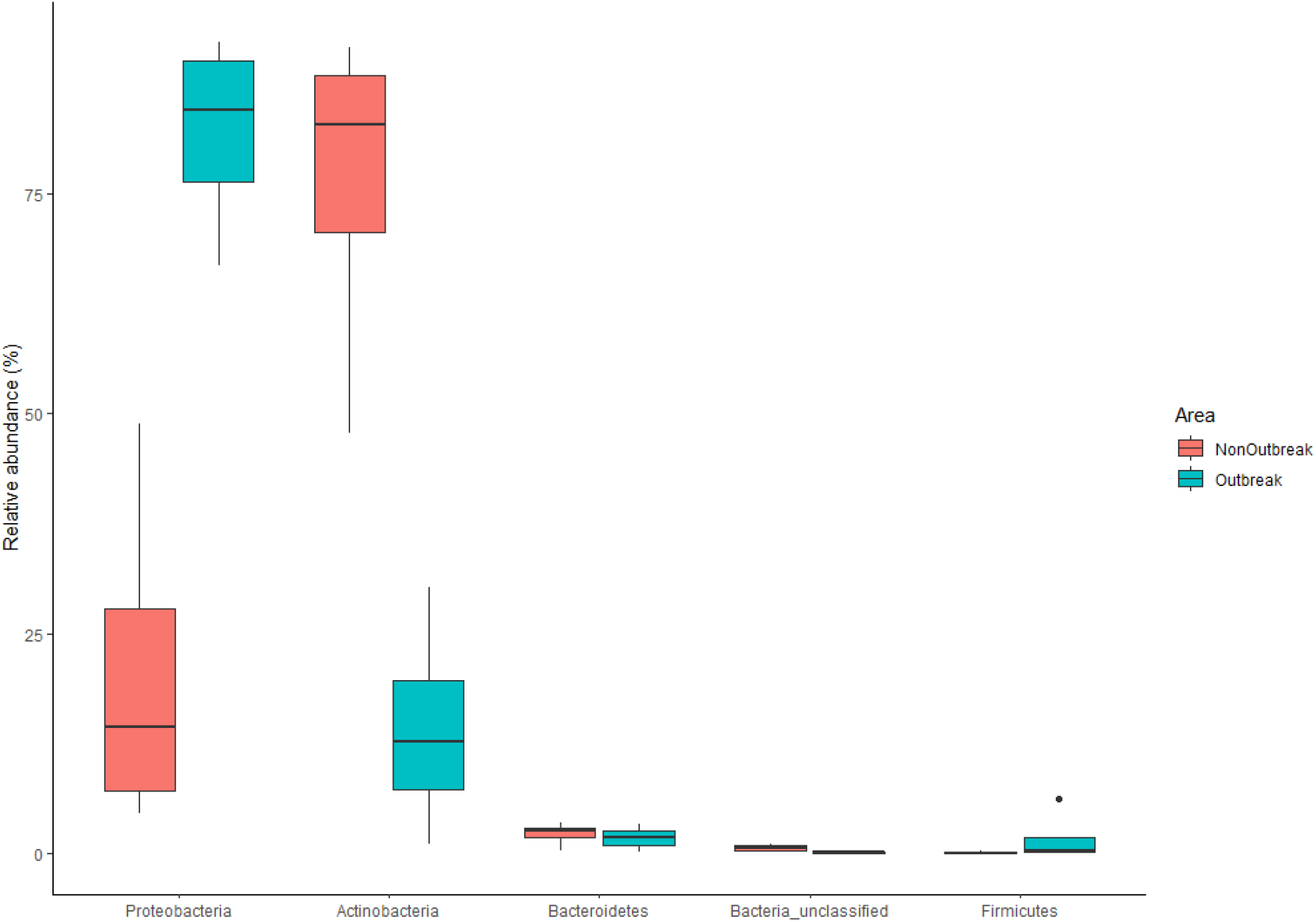
Top 5 relatively abundant bacterial phyla of *M. plana* bagworm larvae in the comparison between non-outbreak area and outbreak area.

The most abundant families consisted of *Enterobacteriaceae*, followed by *Microbacteriaceae, Pseudomonadaceae, Burkholderiaceae, Sphingobacteriaceae, Kineosporiaceae* and other families individually having less than 1% relative abundance (Figure 8 and Table 10). We again compared the relative abundance of families between the 2 areas and found that there were a few families that were significant difference in relative abundance (Table 11). *Enterobacteriaceae* was more abundant in the outbreak area (75.41%) compared to non-outbreak area (11.67%). However, there were more of *Microbacteriaceae* (70.87%) and unclassified bacteria (0.60%) in non-outbreak area compared to outbreak area (12.47% and 0.14% respectively. There were presence of *P3OB-42* (0.10%) and unclassified *Alphaproteobacteria* (0.06%) in non-outbreak area while there were none of the 2 families in the outbreak area.

**Table 10.** Bacterial families with an overall abundance of more than 1% in the comparison between non-outbreak area and outbreak area.

| Families | Non-outbreak area (%) | Outbreak area (%) | Overall Presence >1% (%) |
| --- | --- | --- | --- |
| *Enterobacteriaceae* | 11.67 | 75.41 | 43.54 |
| *Microbacteriaceae* | 70.87 | 12.47 | 41.67 |
| *Pseudomonadaceae* | 5.03 | 3.34 | 4.18 |
| *Burkholderiaceae* | 2.93 | 2.56 | 2.74 |
| *Sphingobacteriaceae* | 1.57 | 1.24 | 1.40 |
| *Kineosporiaceae* | 2.33 | 0.15 | 1.24 |

**Table 11.** Bacterial families with significant difference in relative abundance between non-outbreak area and outbreak area.

| Families | Non-outbreak area (%) | Outbreak area (%) | Overall Presence (%) |
| --- | --- | --- | --- |
| *Enterobacteriaceae* | 11.67 | 75.41 | 43.54 |
| *Microbacteriaceae* | 70.87 | 12.47 | 41.67 |
| Unclassified bacteria | 0.60 | 0.14 | 0.37 |
| *Sphingomonadaceae* | 0.12 | 0.01 | 0.07 |
| *P3OB-42* | 0.10 | 0.00 | 0.05 |
| Unclassified α-proteobacteria | 0.06 | 0.00 | 0.03 |

**Figure 8.**
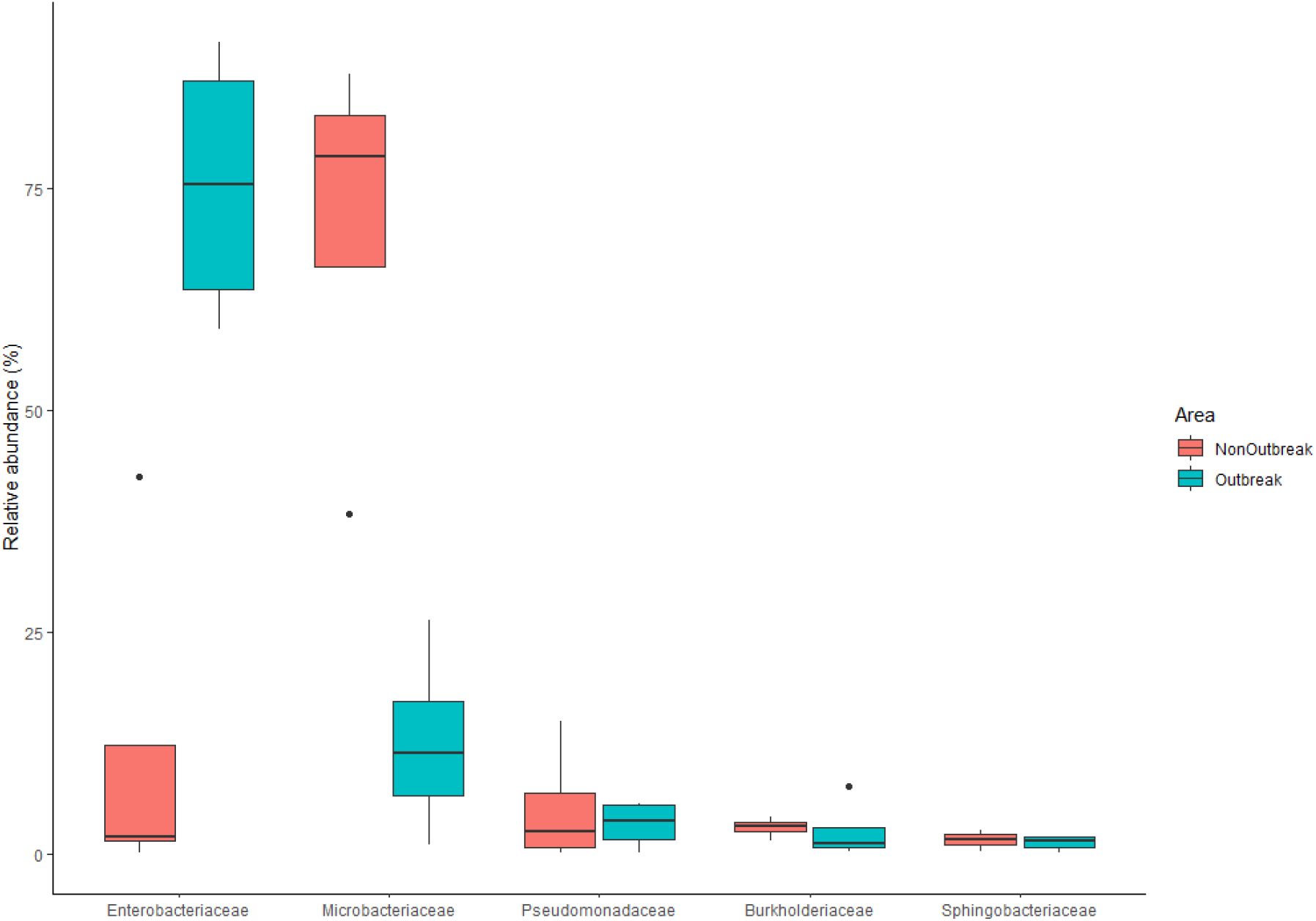
Top 5 relatively abundant bacterial families of *M. plana* bagworm larvae in the comparison between non-outbreak area and outbreak area.

#### Diversity of the bacterial community

The Shannon diversity index showed that the bacterial community from the non-outbreak area had a higher diversity than the counterpart (Figure 9 and Table 12).The number of OTUs was higher in non-outbreak area than outbreak area, revealing that the bacterial community in the non-outbreak area was richer. Shannon evenness was calculated and it showed that the bacterial community in outbreak area was more even compared to the non-outbreak area. However, the diversity, richness and eveness between the non-outbreak area and outbreak area were not significantly different.

**Table 12.** Shannon diversity index, number of OTUs and Shannon evenness of bacterial community in the non-outbreak and outbreak area.

| Area | Sample | Shannon | OTUs | Evenness |
| --- | --- | --- | --- | --- |
| **Non-Outbreak** | NLS0 | 1.691 | 133.576 | 0.345 |
|  | NLS7 | 1.947 | 88.000 | 0.435 |
|  | NLS12 | 1.248 | 92.506 | 0.276 |
|  | NLS16 | 1.093 | 101.029 | 0.237 |
|  | **Average** | **1.494** | **103.778** | **0.323** |
| **Outbreak** | OLS3 | 0.678 | 50.042 | 0.173 |
|  | OLS4 | 1.773 | 104.775 | 0.381 |
|  | OLS5 | 1.893 | 87.202 | 0.424 |
|  | OLS6 | 1.156 | 89.667 | 0.257 |
|  | **Average** | **1.375** | **82.922** | **0.309** |
| **T.Test** | **p-Value** | **0.376** | **0.204** | **0.423** |

**Figure 9.**
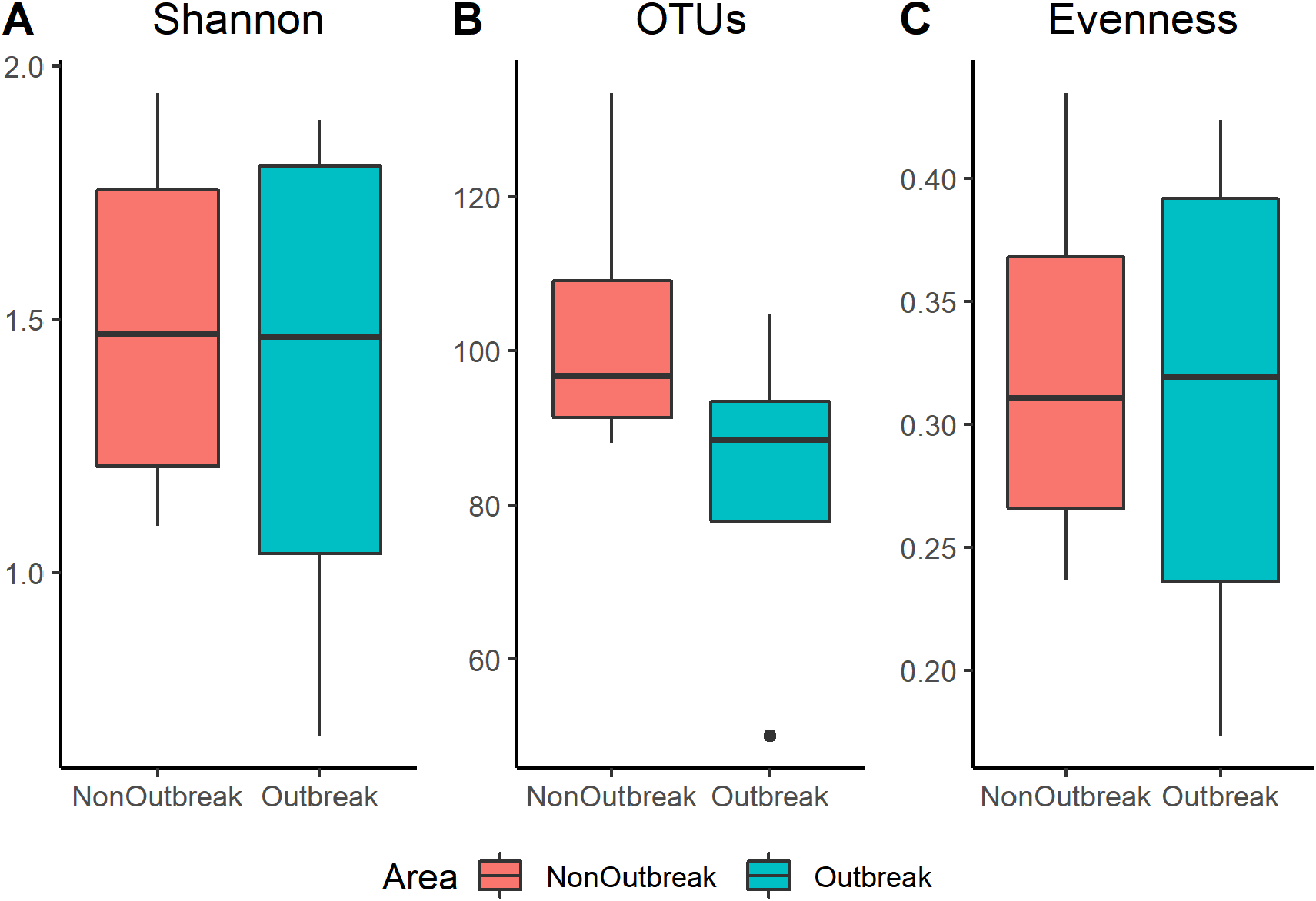
Alpha-diversity of larvae of *M. plana* in the comparision between non-outbreak area and outbreak area. **A**: Shannon diversity index; **B:** Number of OTUs; **C: D:** Shannon Evenness.

**Figure 10.**
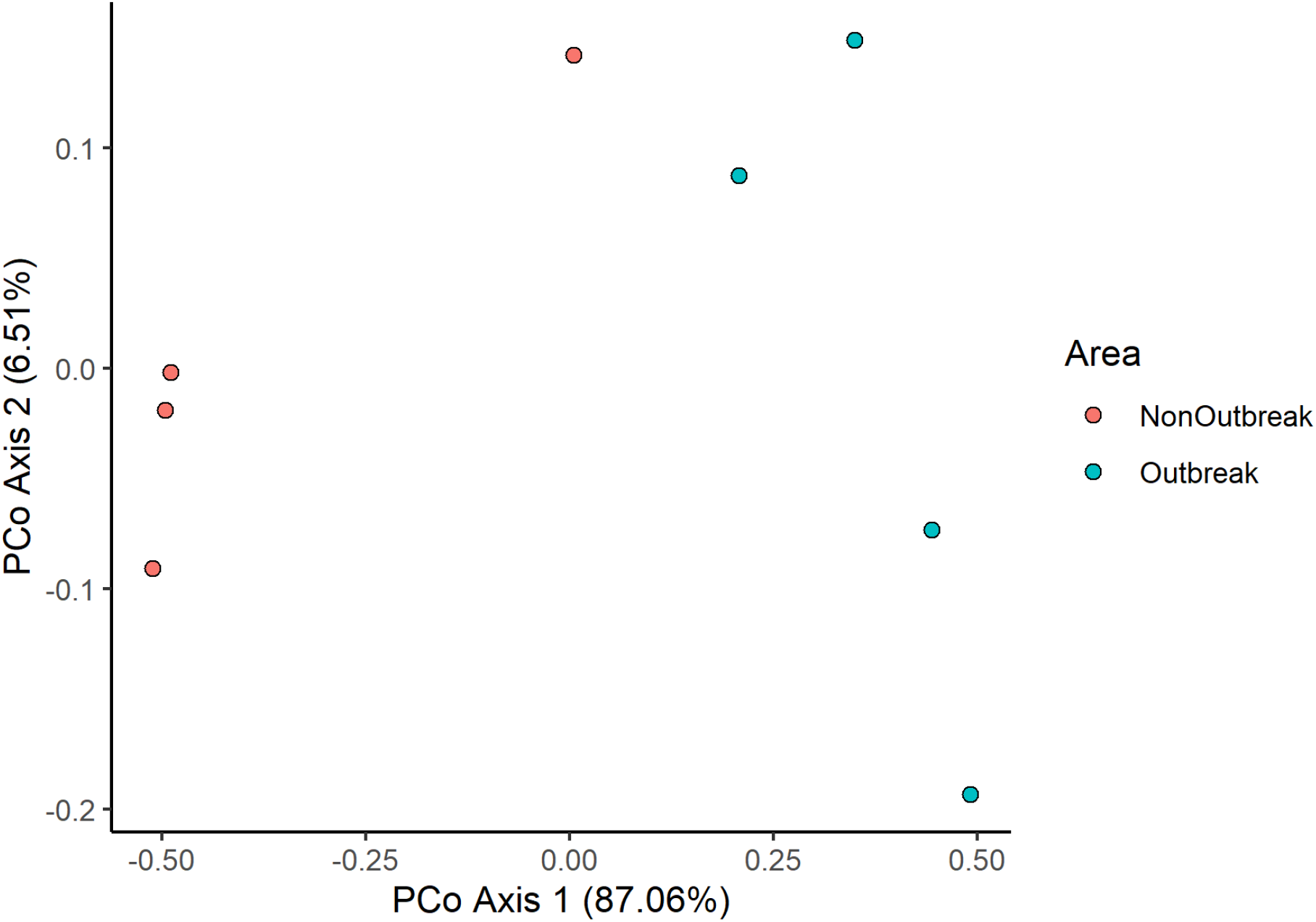
Principal Coordinate Analysis (PCOA) plot of bacterial communities of *M. plana* bagworm larvae in the comparison between areas.

From the PCoA (Figure10), we observed a clear separation between the samples from non-outbreak area and outbreak area. AMOVA test was done on the samples and the result (Table 13) showed separation between the two areas was significantly different. This meant that the bacterial community was different from one another.

**Table 13.** AMOVA test done on samples from non-outbreak and outbreak area. (Significance at p-value < 0.05)

| Non-Outbreak -<br>Outbreak | Among | Within | Total |
| --- | --- | --- | --- |
| SS | 1.087 | 0.269 | 1.357 |
| df | 1 | 6 | 7 |
| MS | 1.087 | 0.045 |  |
| Fs: | 24.209 |  |  |
| p-value: 0.034* |  |  |  |

The HOMOVA test (Table 14) showed that there was a significant difference in the variation of bacterial community between the 2 areas. The non-outbreak area has a higher variation (0.063) compared to the outbreak area (0.027). This showed that bacterial community in the non-outbreak area was less stable than the outbreak area.

**Table 14.**
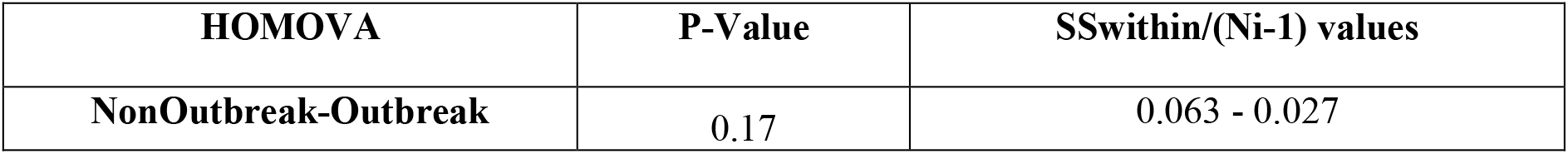
HOMOVA test done on the samples from non-outbreak and outbreak area. (Significance at p-value < 0.05)

## Discussion

### Overview of the Bacterial Community

At present, most studies on lepidopteran microbiota were focused on the microorganisms linked to the larval gut. However, this only provide a smaller but more focused view of the community. In this study, focus was made on the early instar stage as well as the late instar stage from the outbreak area to see whether there was any difference between them. The authors further compared the bacterial community of the late instar stage *M. plana* between non-outbreak area and the outbreak area to investigate spatial-associated shift in the bacterial community. Generally, the bacterial community was dominated by *Proteobacteria* and *Actinobacteria*. These phyla were commonly found within the Lepidoptera order (13,14,30). On the family level, there were a few dominant families such as the *Enterobacteriaceae* and *Microbacteriaceae*. Again, these families can also be found in different Lepidopteran species (13,31,32).

The overall dominance of the specific bacteria in this study may have some sort of beneficial roles to the health of the larvae. Although *M. plana* larvae is polyphagous, they were collected from the foliage of the oil palm tree, and might only have that host plant as its diet. However, it could also be that the larvae obtained these bacteria solely from their environment or diet but provided little or no benefit. Our hypotheses are similar with a study done by Phalnikar and collegeaus (33). In their study, they observed that their most common and abundant OTUs in butterflies were also common in different insect-associated microbiomes.

This lead them to hyphothesise that the insect-bacterial co-occurrencce may indicate evolved functional relationships or it could merely act as ecological or dietary roles. The latter hypothesis might be due to absence or presence of very little resident bacteria found in caterpillar such as in a study done by Hammer and colleageus (34) and is in agreement with Phalniar and colleagues’ study where they found a substantial overlap of bacterial communities from larval and dietary resources which indicated that bacterial communities in larval are mainly influenced by passive procurement of bacteria from dietary resources (33). Nevertheless, it is important to note that the microbiome varies greatly across lepidopteran species and even within species (35). The entire larvae were sampled, yet there was no trace as to where exactly these bacteria reside, although some studies had found that the bacterial communities from the whole insect can be similar to the bacterial communities sampled from the gut (16,36,37).

### Comparison Between Early Instar Stage and Late Instar Stage

Here in this part, we compared the bacterial communities in early instar and late instar stages of *M. plana* from outbreak area. In holometabolous insect such as the bagworm, the bacterial community is affected by their developmental stage (13,38,39). Findings from this study reveals that, the *Proteobacteria* phylum was the most dominant phylum, followed by *Actinobacteria, Bacteroidetes, Firmicutes* and other minor phyla. This is in keeping with a review done by Voiral *et al*. (13) which screened independent studies of 30 different lepidopteran species, and found that *Proteobacteria* phylum was the most common phylum. Another study done on the moth *Brithys crini* at different developmental stages also found the *Proteobacteria* as the most abundant phyla (1). At family level, the *Enterobacteriaceae* was the dominant family and is also commonly found in other Lepideptoran species (13,30,40). However, the bacteria that were significantly different in relative abundance were all from the minor families. This observation could be attributed to the change in the feeding behaviour of the larvae. In the early instar stage, the larvae scrape on the leaf epidermis using their mandibles but changed to cutting leaves at late instar stage (2). The less active feeding behaviour of the larvae at late instar stage could also played a part in this difference in relative abundance of the mentioned families. In a study done by Kok and colleagues (3), they observed that the lab-reared *M. plana* larvae reduced their feeding activities and remained in their cases after the 4^th^ instar stage. If the bulk of the bacterial community of the larvae is obtained from their diet, this reduced feeding activity could have impacted the abundance of certain bacterial species. However, we could only assume that the wild *M. plana* larvae exhibit the same feeding behaviour as the lab-reared larvae. We also hypothesised that the difference in relative abundance of the mentioned minor families might be due to the developmental time from one instar to another. The early instar may be exposed to environmental bacteria for a longer time due to a longer time needed to develop in the early instar stage than in the late stage (3).

In the current study, it was observed that the bacterial diversity, richness, and evenness in the early instar stage were higher than that of the late instar stage. However, the results were not significantly different between the two larval instar stage. It was further shown that the bacterial community structures from early instar stage and late instar stage did not form significantly separated clusters. This observation hinted that the instar stage did not significantly contribute to variability of the bacterial community. In some Lepidopteran species such as *Plodia interpunctella* and *Plutella xylostellai*, their bacterial community did not change across developmental stages (41–43). This could happen to the *M. plana* larvae where its bacterial community structure is not significantly affected by the developmental stage. The similarity in the bacterial community could also be attributed to the larvae having the same host plant (oil palm tree *Elaeis guineensis*), as different diet might influence bacterial communities in different ways such as promoting differential bacterial growth (44–46). The similarity of the bacterial community could also be due to the same area where the bagworms were collected as different environments could affect the microbial community in insect (41,47).

### Comparison Between Non-Outbreak Area and Outbreak Area

Here the authors compared the bacterial community of the late instar larvae from non-outbreak area as well as outbreak area. Similar to the previous comparison, it was observed that the *Proteobacteria* was the most dominant phylum while the *Enterobacteriacea* () was found to be the most common family. The study further revealed some phyla and families that had significant difference in relative abundance between the two areas. Although the bacterial diversity, richness, and evenness between the two areas were not significantly different, there was a clear and significant separation in the centre of the samples clusterwhich implied a significantly different bacterial community structure. Although the bagworms were collected from oil palm plantations and have the same host plant in both non-outbreak area and outbreak area, there could be an underlying environmental factor that attribute to these significant difference. The authors suspect that the bacterial community on the host plant itself or the surrounding environment were different between the non-outbreak area and outbreak area, hence contributing to the difference in relative abundance of the phyla as seen. This phenomenon was also seen in a study done by Jones and colleages (32) where they found distinct bacterial communities in corn earworm midgut from different sites but with the same host plant. In addition, the soil microbiome in the two areas might be different. A study showed that insects that feed on foliar obtained their microbiomes from the soil (48). The authors in the mentioned study stated that the microbiome of the caterpillar that fed on intact plant had a more distinct microbiome and the microbiome resembled the soil microbiomes. In another study (49), the authors found that the caterpillar’s bacterial communities resembled the local soil microbiomes in which the host plant was growing. Their studies provide us with the hypothesis that although the bacterial communities of the bagworm larvae from outbreak area and non-outbreak area generally are similar, the significantly different in abundance of certain bacteria species found in this comparison could be reflected in the difference of local soil microbiome where the bagworms were collected.

## Conclusion

The bacterial commmunities in the larvae of *M. plana* in their early and late instar stages as well as form the non-outbreak and outbreak areas were compared. Although the bacterial communities in the comparisons were not significantly different in terms of diversity, richness and evenness, there were some significant difference in abundane of certain bacteria phyla and families. A significant and clear distinction in the bacterial community structure when comparing non-outbreak area and outbreak area was also recorded. This study provides a first insight to the bacterial community of the *M. plana* larvae. However, more studies are needed to uncover the bacterial communities in greater details especially the gut microbiome.

## Acknowledgements

We would like to thank the team from the Entomology Research Group, Pest Management Unit, FGV R&D Sdn Bhd, Felda Jengka 7 and Felda Gunung Besout 02/03 for providing study sites and for the help of the team to conduct sampling of bagwoms samples in the field.We would also like to thank Ahmad Mustakim bin Mohamad, Ahmad Faisal, Mardani Abdul Halim and Azali Azlan for their advices in the statistical and molecular part of the work.

## Author Contributions

Andrew Chung Jie TING contributed to the conceptualization, methodology, formal analysis, investigation, visualization, writing (original draft), and writing (review and editing) of the project. Cik Mohd Rizuan ZAINAL ABIDIN contributed to the conceptualization, methodology and investigation, resources of the project. Noor Hisham HAMID contributed to the conceptualization, methodology and investigation, resources of the project. Ghows AZZAM contributed to the conceptualization, methodology, resources, writing (review and editing), supervision, project administration, and funding acquisition of the research. Hasber SALIM contributed to the conceptualization, methodology, resources, writing (review and editing), supervision, project administration, and funding acquisition of the research.

